# Open-hardware solutions for light sheet microscopy specimen chambers

**DOI:** 10.1101/2024.08.22.609188

**Authors:** Artemiy Golden, Julien Colombelli, Ernst H.K. Stelzer, Francesco Pampaloni

## Abstract

Light sheet fluorescence microscopy (LSFM) is a powerful tool for imaging large three-dimensional biological samples. However, the design and fabrication of specimen chambers for these systems present significant challenges, particularly in maintaining water-tight seals, preventing contamination, and ensuring the flexibility needed for precise positioning of the objective and sample. This study introduces open-hardware solutions to address these challenges, utilising a combination of 3D printing, silicone injection moulding, and FEP-foil thermoforming. We describe the development of custom, highly flexible silicone seals and connectors through a laboratory-scale injection moulding process. These components enable precise, low-resistance movement of imaging objectives and specimen holders, which is crucial for maintaining imaging accuracy. Additionally, we introduce a novel “optical window” design that isolates the objective lens from the immersion medium, significantly reducing the risk of contamination and facilitating easy exchange of chambers and lenses without compromising sterility. The practicality of these designs is demonstrated through their application in long-term live imaging of *Tribolium castaneum* embryos, honey bee embryos, and human mesenchymal stem cell spheroids. By providing open-source CAD and 3D printing files, this work promotes accessibility and customization in microscopy, enabling researchers to easily replicate and adapt these solutions to their specific needs.

## Introduction

Light sheet fluorescence microscopy (LSFM) is the preferred technique for imaging large three-dimensional samples, such as cellular spheroids, organoids, and embryos. Two decades after the introduction of selective plane illumination microscopy (Single Plane Illumination Microscopy, SPIM) by Huisken in 2004 (1), and Keller 2008 (2) (Digitally Scanned Light Sheet Microscope, DSLM), several LSFM implementations have emerged, including single objective lens systems compatible with conventional sample holders like multi-well plates. Nevertheless, the classical SPIM setup, featuring a central imaging chamber in which both the sample and the detection lens are immersed in an index-matching liquid remains, with variations (3), widely used (**Figure 1**).

**Figure 1.**
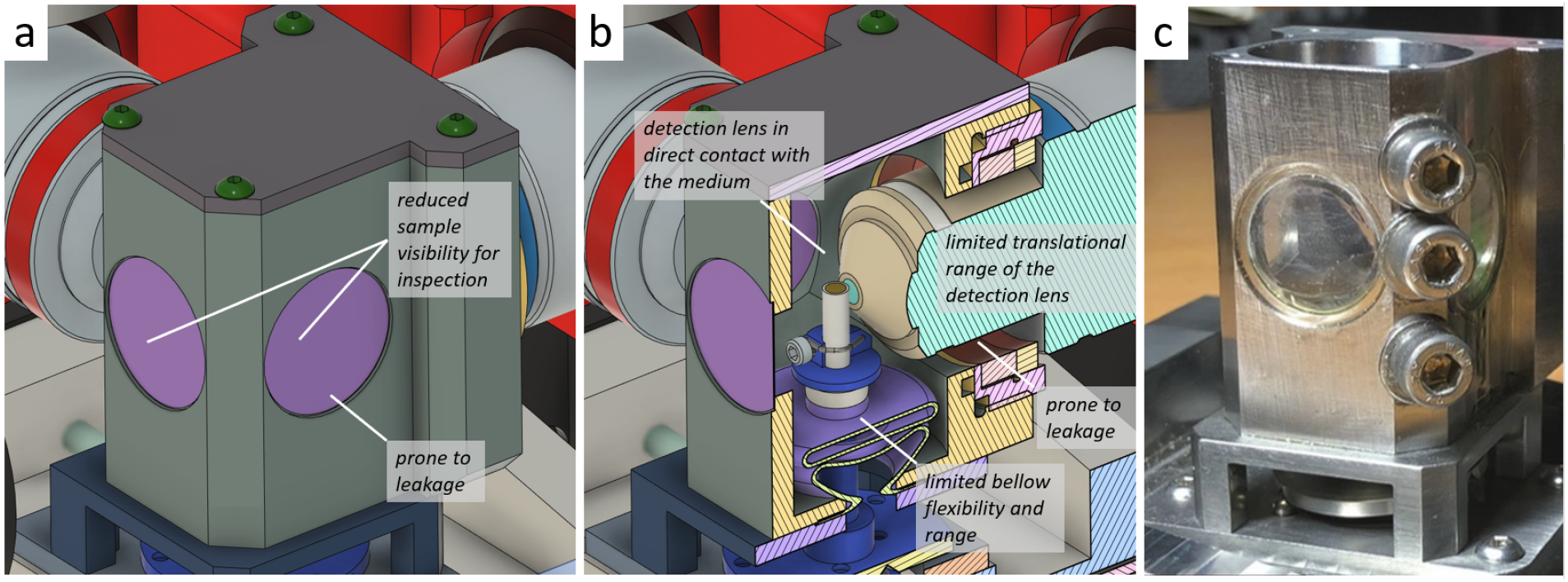
Design challenges of a conventional SPIM-like LSFM chamber. (a) Chamber from outside: off-the-shelf 16 mm round glass coverslips as inspection windows are tiny and make sample centering and positioning troublesome. Moreover, glueing the coverslip to the chamber creates a leakage-prone interface. (b) Chamber inner structure: During long-term live imaging, the detection objective lens is plugged into the chamber in direct contact with the immersion medium, increasing the risk of bacterial/fungal contamination. Also, the medium itself can deteriorate the lens. Off-the-shelf sealings limit the axial translation of the lens and are prone to leakages due to fast ageing. Moreover, off-the-shelf rubber bellows do not allow for a highly flexible water-tight connection of the specimen holder. (c) A photograph of a real chamber, machined in stainless steel.

Indeed, from a practical point of view, the design of a SPIM imaging chamber presents the biggest challenges to LSFM-builders, given several limitations posed by this configuration: 1) The translation range of the imaging lens is limited due to rigid seals. The detection lens seal must be easy to install, reliable, and flexible enough to provide low resistance if the objective lens is supposed to translate along its optical axis. 2) The objective lens is in direct contact with the medium, which can lead to bacterial contamination. Conversely, the medium can contaminate the objective lens with some toxins/drugs used in the experiment. However, designs allowing the insulation of the objective lens from the immersion medium are lacking so far. 3) To minimise potential leakage points, the assembly of the chamber should be optimised. Screwed elements are particularly prone to leaks at their interfaces. 4) Unplugging the chamber without emptying the immersion medium is not possible. Yet, this feature would be highly desirable for a fast and safe specimen turnover, especially for samples that need to be further cultured and possibly re-imaged after a certain time interval. 5) In general, the chamber must be designed for easy and cost-effective manufacturing to ensure the availability of multiple chambers when using different imaging media or handling toxic substances.

Additionally, for configurations where the specimen holder is mounted on the bottom of the specimen chamber (like in the mDSLM (4)) there are further design requirements for a sealing solution. It needs to allow for the holder movement in X, Y, Z directions and rotation, while maintaining a watertight seal and not applying significant force on the holder to prevent deflection.

Overall, the challenging design aspects of the imaging chamber have heavily impaired the user-friendliness of SPIM-like implementations (**Figure 1**). To address these challenges, developers of custom light-sheet systems are increasingly turning to 3D-printed specimen chambers (5–7). 3D printing allows for easy customization and manufacture. Many readily available plastic materials make the chambers biocompatible(8,9) and inexpensive to manufacture with technologies such as FDM (fused deposition modelling) and SLA (stereolithography). However, methods for the design and fabrication of highly flexible water-tight seals and connections are still lacking. Additionally, to the best of our knowledge, methods for the quick exchange of the sample chamber or a solution for insulating the imaging objective lens from the immersion liquid have not been described before.

We present an open hardware approach to the design and manufacture of specimen chambers addressing the challenges mentioned above. We first show how to fabricate highly flexible custom seals for objectives and specimen holders. Next, we provide innovative solutions based on an “optical window” compartmentalization to prevent media contamination by plugged objective lenses and to allow for a straightforward chamber exchange. Finally, we showcase specimen chamber implementations for different SPIM-derived light-sheet setups and include CAD and 3D printer project files to easily reproduce our designs. One of the designs allowed for the first long-term live imaging of a honeybee development demonstrating how even unusual engineering requirements can be met by our design and manufacturing approach.

## Results

### 1. Fabrication of flexible seals and bellows with injection moulding

For the fabrication of mechanically highly flexible elements a laboratory-scale injection moulding process was developed. In the process, a two-component curable and biocompatible silicone elastomer is pre-mixed and injected in a 3D-printed mould by using a common syringe inserted in a pre-defined inlet. The mould components are tightly and precisely assembled by using threaded parts and screws (**Figure 2**). With this approach, several highly flexible elements were fabricated: 1) thin open diaphragm water-tight seals for objective lenses (**Figure 2a**), 2) Bellows allowing large XYZ displacements plus rotation of the sample holder (**Figure 2b**), and low-force connectors for large displacement of objective lenses (**Figure 2c**). The design and practical details of the injection moulding process, as well as suggestions on how to avoid the formation of bubbles, are described in the *Material and Methods* section and the supplementary material, whereas their applications are showcased in the following *Results* sections.

**Figure 2.**
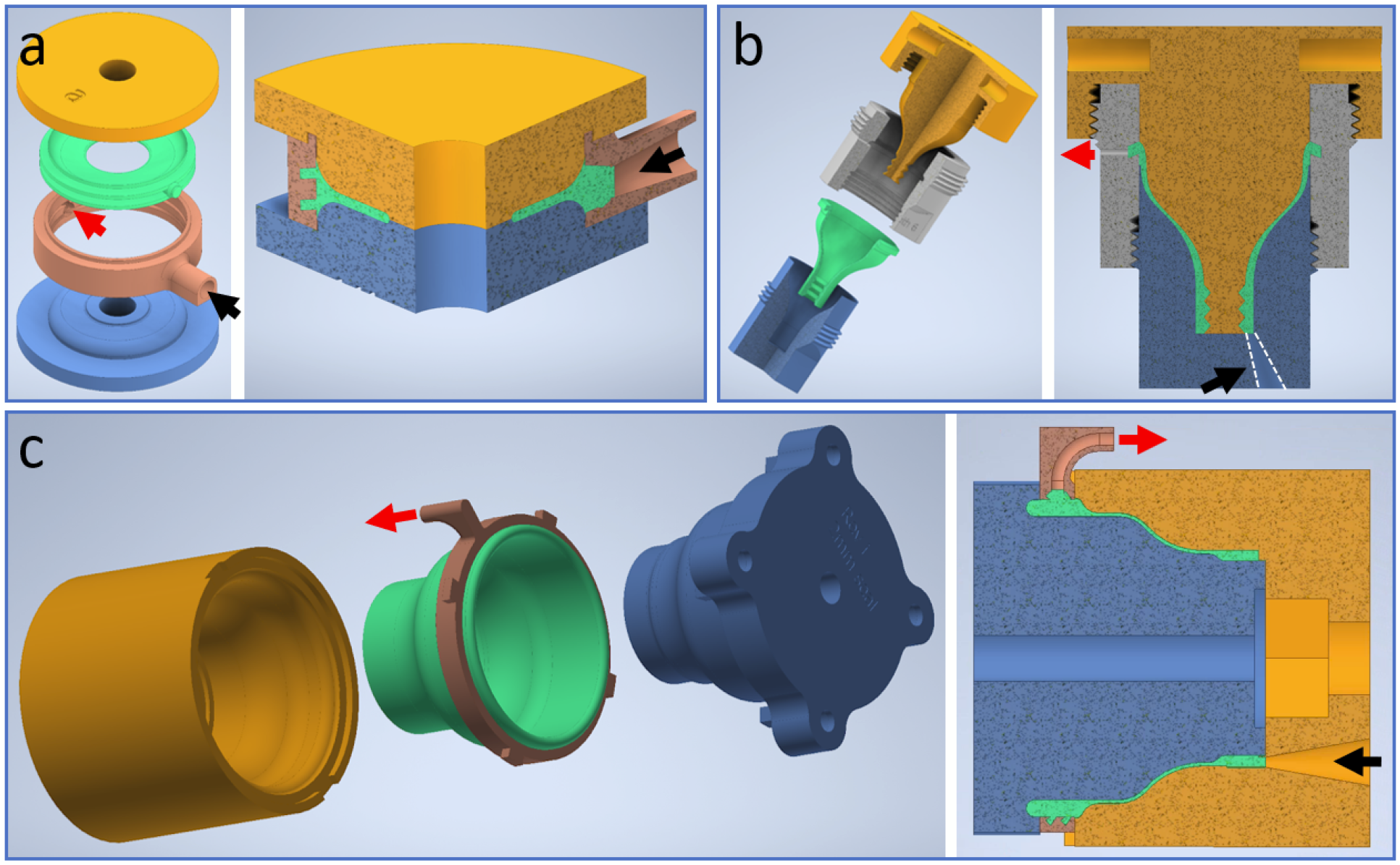
Fabrication of highly flexible connectors for an LSFM chamber. (a) Mould system for the **membrane objective lens seal**. The mould consists of three main components: two primary components (blue and yellow) that define the shape of the silicone membrane (green), and a third component which is the frame to which the open diaphragm is anchored. The frame allows one to mechanically connect the membrane to the chamber body. The primary components are secured together with a screw (not shown) passing through a central hole. The black arrow indicates the inlet for the silicone rubber injection and the red arrow the outlet. (b) Mould system for the **sample holder bellow**. In this case, the two primary components (blue and yellow parts) are tightened together by a central joint (grey). The inlet is indicated by the black arrow, and the outlet by the red arrow. The silicone bellow is shown in green. (c) **Low-force objective connection**. The two primary components (blue and yellow parts) are tightened with screws (not shown). The resulting silicone part (green) is anchored to a bayonet system (brown) to connect with the LSFM chamber. The inlet for the silicone rubber injection is indicated by the black arrow and the outlet by the red arrow.

### 2. Open diaphragm water-tight seal for objective lenses

LSFM sample chambers require water-tight seals around the objective lens. These seals must be sufficiently flexible to accommodate the translation of the objective lens along the optical axis, enabling chromatic aberration correction and deep tissue imaging. Conventional elastomeric O-rings, which rely on compression to maintain a tight seal, are unsuitable for this purpose as they hinder the necessary movement of the objective lens. Using off-the-shelf radial shaft seals (commonly known as “Simmerrings”) in SPIM-like setups has been a practical solution. However, in our experience, the ageing of elastomer materials and the degradation of the stainless-steel circular springs over time rapidly compromise the water-tightness of these seals. The unavailability of off-the-shelf Simmerrings in the exact sizes needed for a specific objective lens is a further drawback. Thus, custom sealing solutions realisable in a laboratory setting are needed.

Open diaphragm seals use a thin, flexible membrane barrier to create a seal and separate an enclosure between a moving and a stationary component. Their flexibility and ability to accommodate movement make them well-suited for situations where a seal must be maintained while allowing translation along an axis, as in the case of an objective lens in LSFM. To achieve a water-tight connection, we followed the rule of thumb of designing an aperture with a diameter that is 50% of the objective lens’s diameter (**Figure 3**).

**Figure 3.**
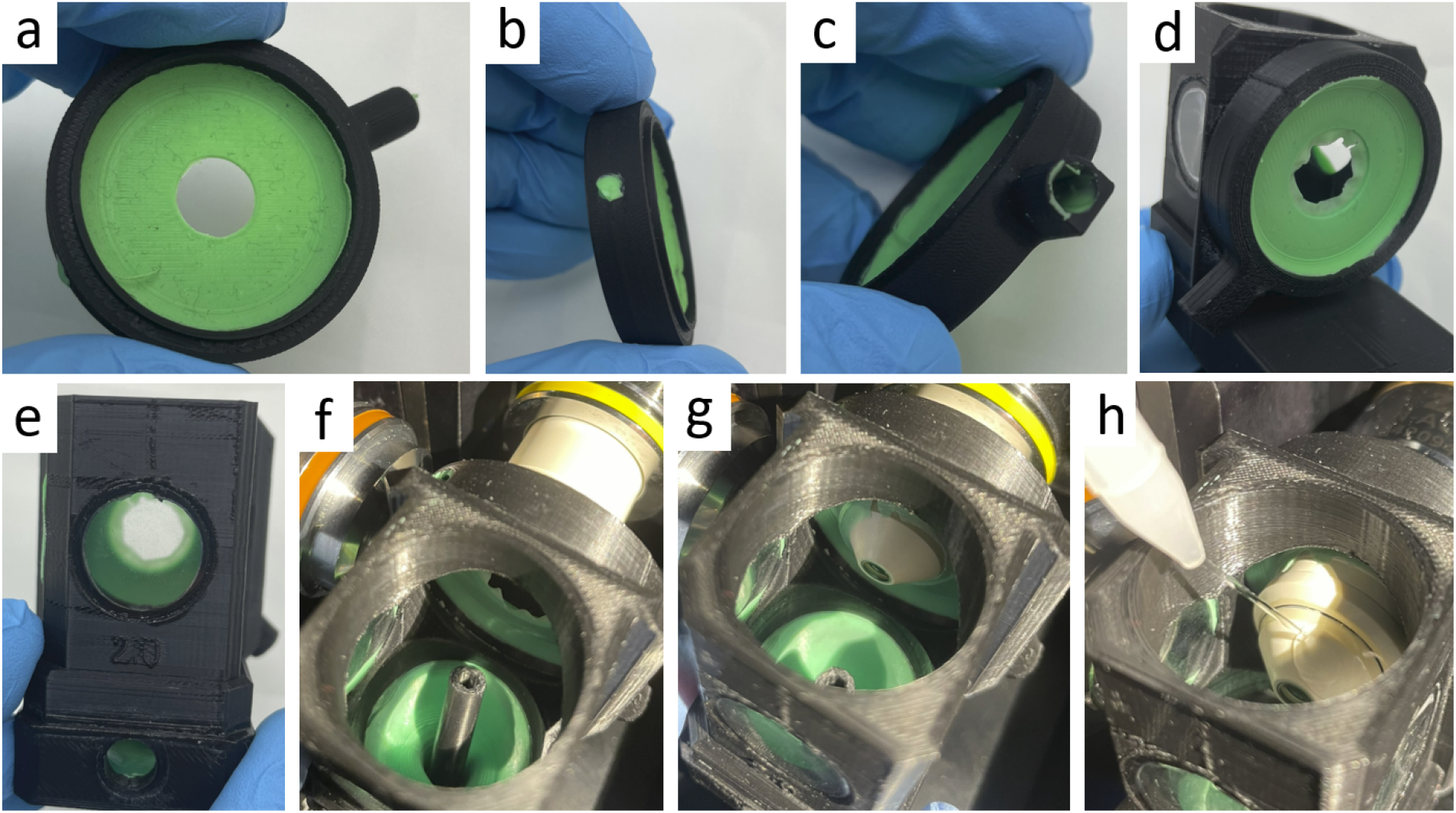
Open diaphragm sheet seal for medium-dipping objective lenses. (a) Front view of the diaphragm seal shown in Figure 2a. (b) Side view of the diaphragm seal showing the silicone rubber injection outlet. (c) Side view of the diaphragm seal showing the silicone rubber injection inlet. (d) diaphragm seal mounted to the LSFM chamber. (e) LSFM chamber, front view. (f-g) Insertion of the LSFM chamber. (h) Filling of the chamber with the immersion/imaging medium.

This approach allows the elastically deformed rim of the diaphragm’s hole to exert pressure on the objective, providing a water-tight yet highly flexible seal (**Figure 4**).

**Figure 4.**
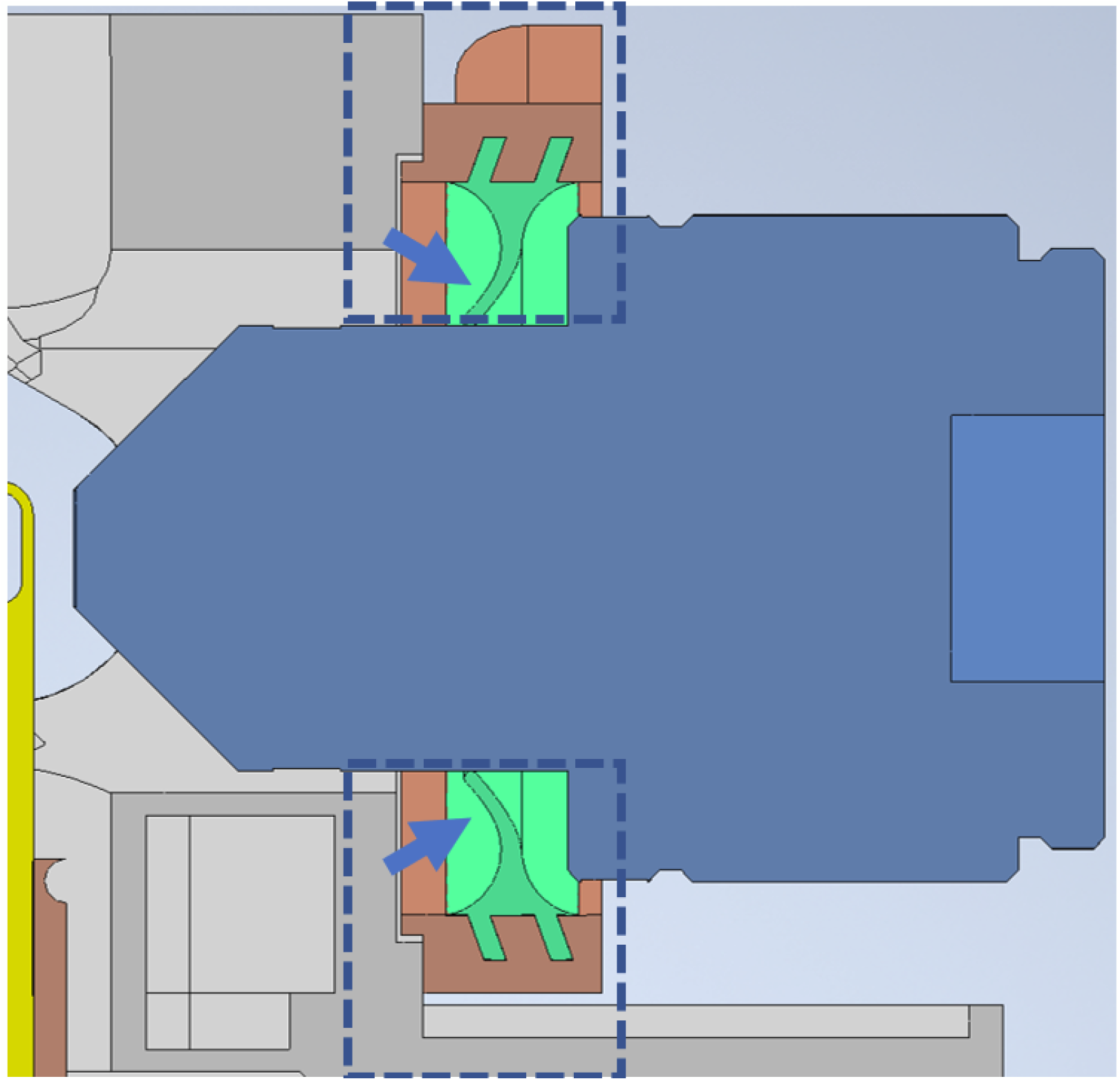
The CAD rendering shows how the open diaphragm creates a water-tight seal by exerting pressure on the objective lens (blue arrows). To achieve this, the diaphragm hole diameter should be roughly 50% smaller than the diameter of the objective lens.

### 3. Low-force connection combined with water-tight bayonet connection for high numerical aperture immersion objectives

In many cases, LSFM setups require a long-range translation of the detection objective lens for focusing and correcting chromatic aberration. This necessitates that connectors and seals for the objective are flexible and exert low force on the objective to avoid interfering with the precise positioning by the linear actuator.

We designed a specimen chamber that fulfills these requirements for a double-detection double-illumination DSLM (dsDSLM) setup. The dsDSLM uses air objectives for illumination and high NA immersion objectives, such as the Plan-Apochromat 20x/1.0W Zeiss objectives, for detection. The detection objectives are mounted on linear actuators for dynamic focus adjustment.

To ease the mounting and sealing of the objectives, as well as the exchange of objectives of different sizes, we developed a bayonet-type locking mechanism for the rapid detachment and release of the lens-seal assembly from the specimen chamber body, where the rubber/silicone moulded seal acts like a spring to ensure a water-tight connection. The specimen chamber design, together with the lens-seal assembly that satisfies all these requirements, is shown in **Figure 5**. Injection moulding of silicone parts allows for a much more simplified design compared to creating such a flexible seal from off-the-shelf parts and 3d printed components only (7).

**Figure 5.**
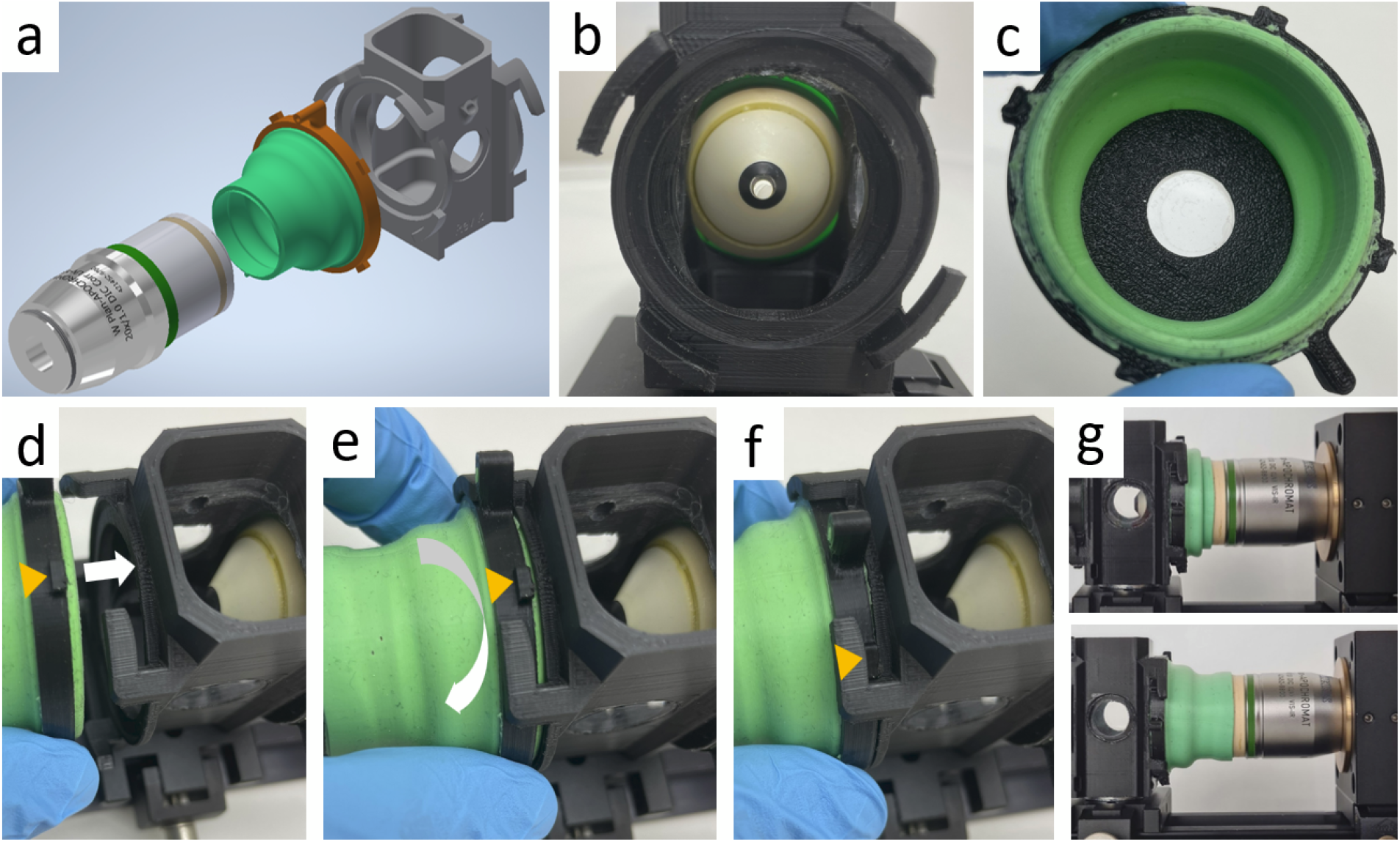
Low resistance sealing solution for large high NA immersion objectives for a dsDSLM setup. (a) a CAD model of an example 20x 1.0 NA Zeiss objective and its sealing solution (Fig. 2c) for a double-sided light-sheet setup. The silicone seal (green), seal holding clamping ring (brown), and the specimen chamber (grey) are shown. The seal ring can be easily clamped using a bayonet-type coupling mechanism that provides a reliable watertight seal. (b) Front view of the water-tight bayonet coupling at the chamber. (c) View of the bayonet coupling mechanism at the flexible seal. (d,e,f) Connection of the LSFM chamber with the flexible seal through the water-tight bayonet coupling. (g) Images of the 20x 1.0 NA objective in the specimen chamber, sealed with the flexible silicone sealing ring clamped to the specimen chamber body. Two positions of the objective show a large range of motion that the silicone seal allows. The silicone seal folds in a predetermined manner allowing for a very low resistance motion of the objective.

### 4. A highly flexible bellow for long-range translation of the sample in the LSFM chamber

The primary advantage of LSFM designs with a SPIM configuration, where the objectives are arranged horizontally and the specimen is inserted vertically, is the ability to image specimens from multiple angles. This capability, combined with the registration and fusion of multiple views, enables the creation of isotropic resolution images, even for thick live samples like *Tribolium castaneum* (3). However, since the specimen holder must rotate within a space constrained by the objective lenses, imaging multiple specimens simultaneously can conveniently be achieved by stacking them vertically on a specimen holder (**Figure 6c**) (10). Vertical stacking requires a large range of motion along the Y-axis of the specimen holder, which must be accommodated by a sealing solution surrounding the holder. While setups with the specimen holder inserted from the top can manage this easily, setups in which the holder is mounted at the bottom demand a flexible seal that allows for extensive, unrestricted movement along the Y-axis, along with sufficient motion along the X and Z axes without exerting significant lateral forces that could cause deflection of the specimen holder.

**Figure 6.**
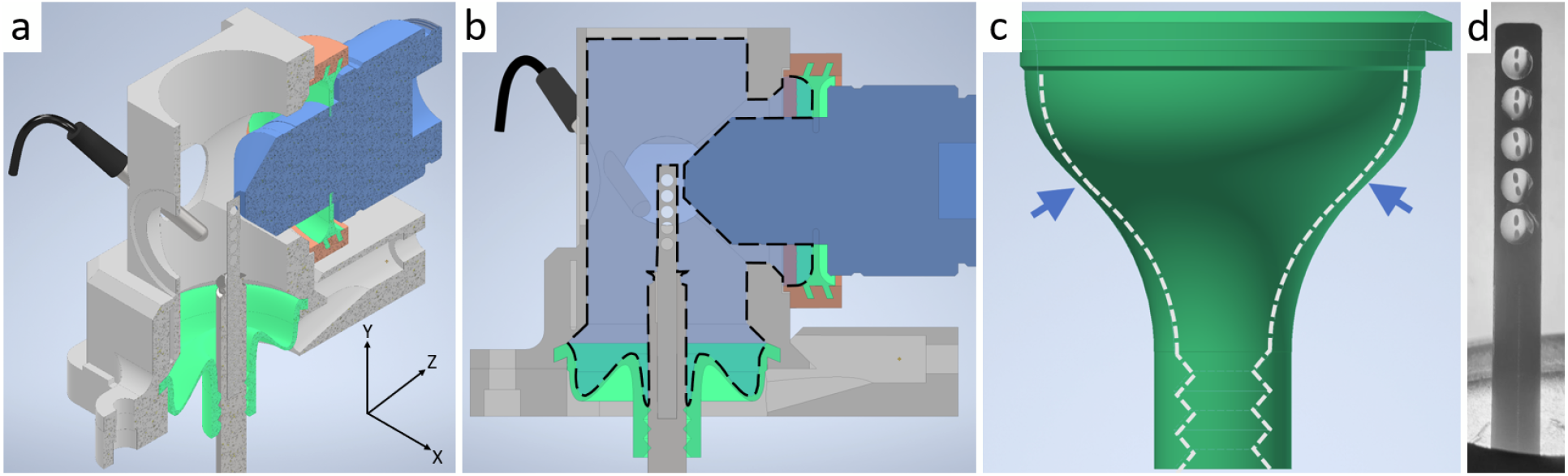
(a,b) CAD renderings of the mDSLM setup specimen chamber showing open diaphragm detection objective sealing solution (Figure 6a) and bellows type sealing (Figure 6b) for the bottom mounted specimen holder. The immersion medium is contained in the volume within the dotted line. The medium is retained at the bottom by the bellow and at the objective lens by the diaphragm. Figure 6c displays the bellows’ wall thickness profile, highlighting the thinnest region (indicated by the arrow), which serves as the flexural surface during the axial translation of the specimen holders. This chamber design allows for specimen positioning with a wide range of motion in X, Y, Z and free rotation. A long range of motion in Y direction (around 20 mm) is especially notable, since it allows for imaging many vertically mounted specimens at once (Figure 6d) and is hard to achieve with other sealing solutions (4,7). The diaphragm-type seal (see also **Figure 3**) provides a very thin profile which is important for spatially constrained environments. The seal is still flexible enough to allow for reliable positioning with PIFOC focus scanners.

To address this, we developed a sealing solution that features a large silicone bellow featuring a variable thickness profile, which allows the silicone to fold with minimal resistance during movements along all three axes (**Figure 6a, b**). The design incorporates the equivalent of several O-rings, ensuring reliable water tightness. To facilitate low-friction rotation of the specimen holder within this silicone seal, we used PTFE-based grease, one of the few lubricants compatible with silicone seals. The repmould used to manufacture this seal is shown in **Figure 2b**. Our design is significantly simpler, requires less parts and is easier to manufacture compared to existing sealing solutions for bottom mounted holders that typically require a separate bushing for a water-tight rotational motion and some form of bellows for translational movements (4,7).

### 5. Optical window for a removable LSFM chamber

A common characteristic of most “classical” chambers for LSFM designs, such as SPIM (Selective Plane Illumination Microscopy) (1), is that the imaging objective lens is in direct contact with a liquid, typically aqueous media, in which the specimen is immersed. This design presents several well-known caveats. Firstly, the objective lens can contaminate the media in various ways. For example, in cell culture media, preventing bacterial contamination is challenging since the lens cannot be autoclaved, and sterilisation with ethanol/water solutions or other disinfectants is not 100% effective, especially during long-term live imaging of 3D cell cultures or organoids over several days (11). Secondly, it is not straightforward to remove or exchange the chamber without first removing the immersion media. Therefore, experiments requiring the imaging of multiple specimens, such as organoids, compromise chamber sterility. Thirdly, in SPIM, there is no simple way to exchange the objective lens during an imaging session, e.g. to achieve multiscale imaging at various magnifications like with conventional inverted or upright microscopes.

The solution we propose to circumvent these limitations is to implement an “optical window” between the imaging objective lens and the chamber’s interior, tailored to the front shape of the dipping lens This approach isolates the LSFM chamber medium from the lens and the external environment, allowing the chamber to remain sealed and sterile. Consequently, the chamber can be easily removed from the microscope without draining, spillage, or contamination of the immersion media. This setup would facilitate serial imaging of multiple samples in separate chambers and allow for the exchange of objective lenses with different magnifications and numerical apertures.

For the optical window to function effectively, it must meet specific criteria. It should have a refractive index close to that of the media in the chamber, e.g. water (n = 1.33), to minimise optical aberrations due to refractive index mismatch, exhibit high transparency in the visible spectral range, be chemically inert and biocompatible, and be mechanically robust as well as thin enough to minimise the reduction of objective lens working distance

Fluorinated polymer foils (FEP foils) have been shown to possess these physical and chemical properties, making them suitable for light microscopy. FEP foils have been extensively used to fabricate sample holders for light sheet microscopy and have been employed to image microtubules, organs, and organoids, both live and optically cleared (12– 15).

Inspired by these previous works, we developed an optical window by vacuum thermoforming FEP foil (14). A suitable positive mould was 3D printed (**Figure 7a**) and used to vacuum form the optical window (**Figure 7b, Figure 8**). The thermoformed optical window features a truncated conical shape to fit the front contour of the objective lens. This approach allows for a straightforward design and fabrication of the optimal shape for each specific type of objective lens. Additionally, more versatile geometries that can accommodate different objective lenses are also feasible.

**Figure 7.**
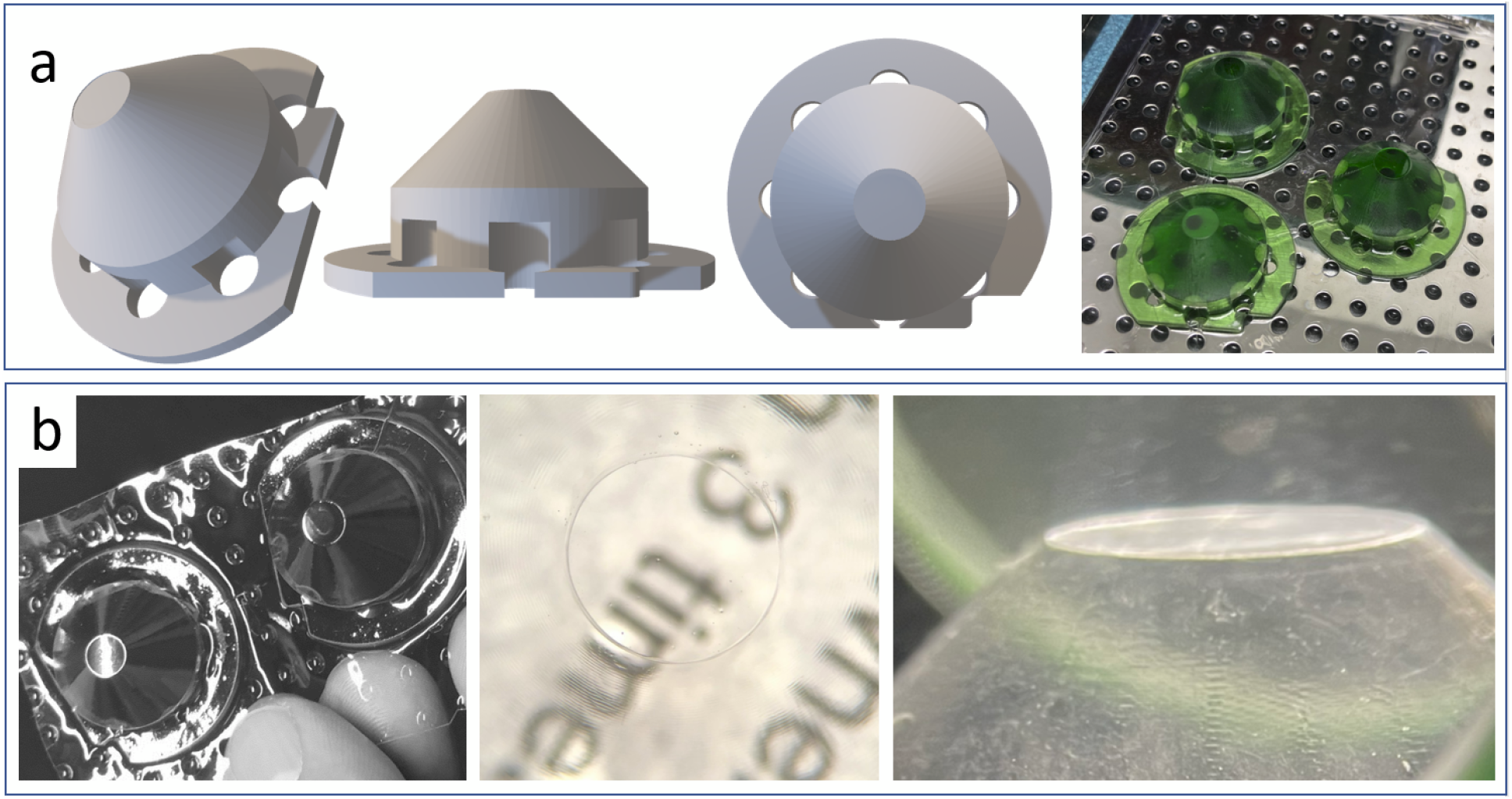
Fabrication of the LSFM chamber optical window. (a) three different views of the positive mould. The mould reproduces the conical shape of the objective lenses’ end. Right panel: photograph of three moulds placed on the thermoforming plate for multiple foil windows production. (b) FEP-foil after thermoforming. Left panel: the mould’s shape has been precisely transferred to the foil. Multiple thermoformed shapes can be produced simultaneously and easily cut out with scissors; middle panel: demonstration of the transparency of the optical window corresponding to the objective’s front lens; right panel: lateral view of the flat FEP-foil window highlighting the flatness of the optical window.

**Figure 8.**
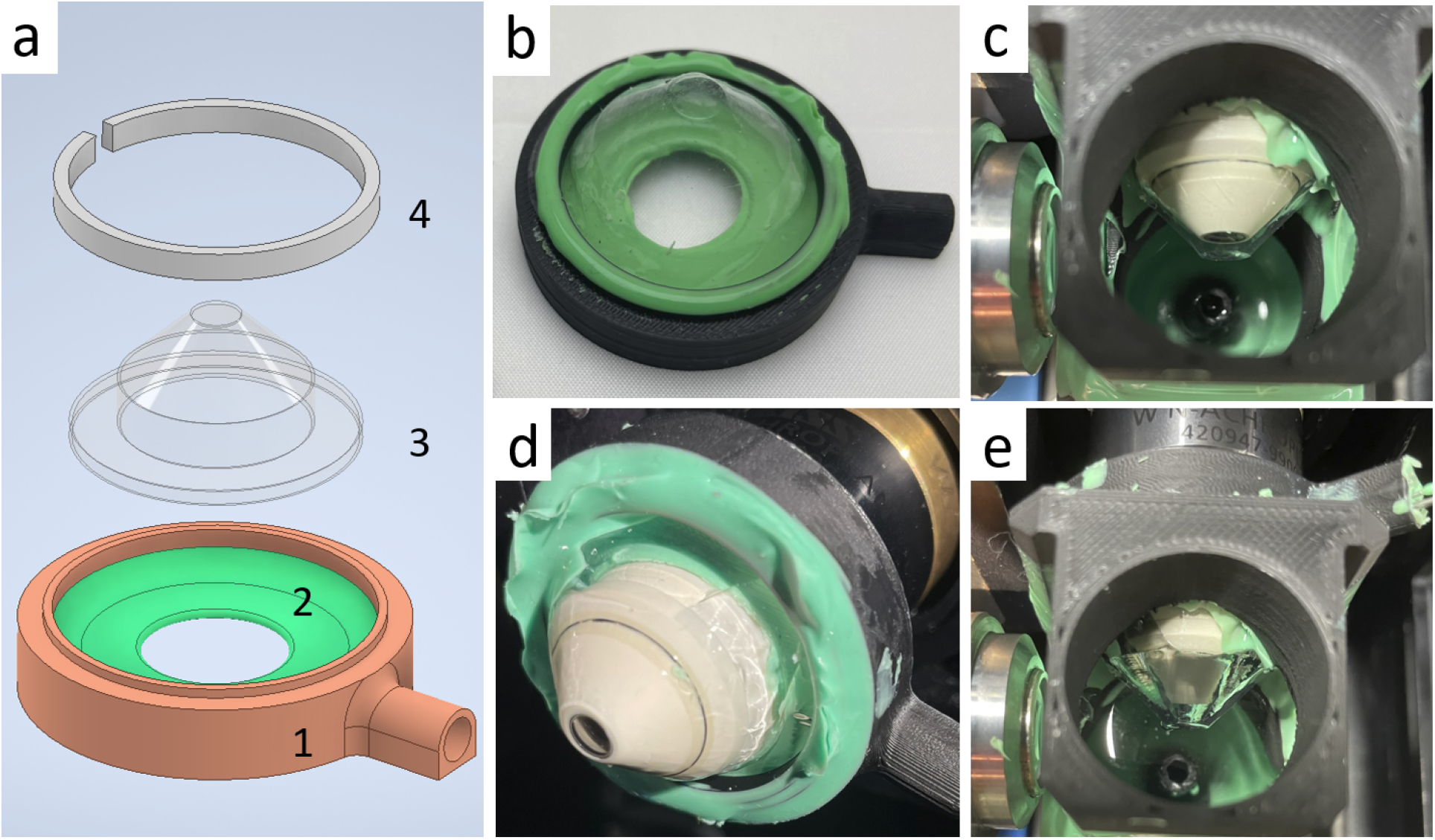
Assembly of the optical window system and integration in the LSFM chamber. (a) The thermoformed optical window (3) is inserted from the top in the open diaphragm (1 + 2) (see also **Figure 3**) and held in place by a counter-ring (3), which is then attached to the diaphragm frame with two-component epoxy glue, ensuring a water-tight connection between the optical window and the adapter membrane. (b) The photograph displays a fully assembled open diaphragm-optical window system. (c) The picture shows the LSFM detection objective lens inserted in the optical window assembly., not connected to the LSFM chamber for illustration purposes. Two cannulas (inlet and outlet, arrow) puncture the silicon rubber and access the gap between the optical window and the objective lens. The outlet allows air expulsion during the insertion of the assembly and the injection of a tiny amount (<100 µl) of immersion media (e.g. water) in the gap between the objective lens and optical window through the inlet. (d) The image shows the optical windows still not completely inserted in the objective lens so that a large gap between the objective’s front lens and the optical window is visible. (e) After completely inserting the optical window, a tiny volume of water is injected into the space between the objective lens and the optical window, filling the gap matching the refractive index. Finally, the LSFM chamber is also filled with the desired aqueous media to start imaging. After imaging, the chamber can straightforwardly be removed without spillage of the inside medium.

The insertion of a new optical interface, such as the FEP-foil window, should have minimal impact on optical quality, introducing only minimal and acceptable aberrations. A primary condition is that the window should be flat, hence the 3D-printed positive mould should exhibit good flatness. Next, the foil should be presented to the lens perpendicularly to its optical axis. To achieve this, a watertight optical window assembly was realised by joining the optical window with the open diaphragm seal described in the previous paragraph 2. In this configuration, an intermediate chamber is obtained between the foil and the front lens, with minimal thickness (50 µm). Herein, a small amount (< 100 µl) of index-matching liquid (e.g., water) can be injected through an inlet (**Figure 8**) to avoid thin air layers. This way, a uniform refractive index is achieved on either side of the foil.

To test the imaging performance of the optical window, cellular spheroids derived from human mesenchymal stem cells (hMSC) were used. The spheroids were fixed with paraformaldehyde, and the nuclei were stained with DAPI. Subsequently, the hMSC spheroids were pipetted into an FEP-foil cuvette for imaging (**Figure 9a**) (14,15) and then placed in the LSFM in front of the optical window (**Figure 9a, right**). A Zeiss W N-Achroplan 10x/0.3 M27 objective lens was used for imaging. Image stacks of the spheroids were recorded with a spacing of 2.6 µm along the z-axis.

**Figure 9.**
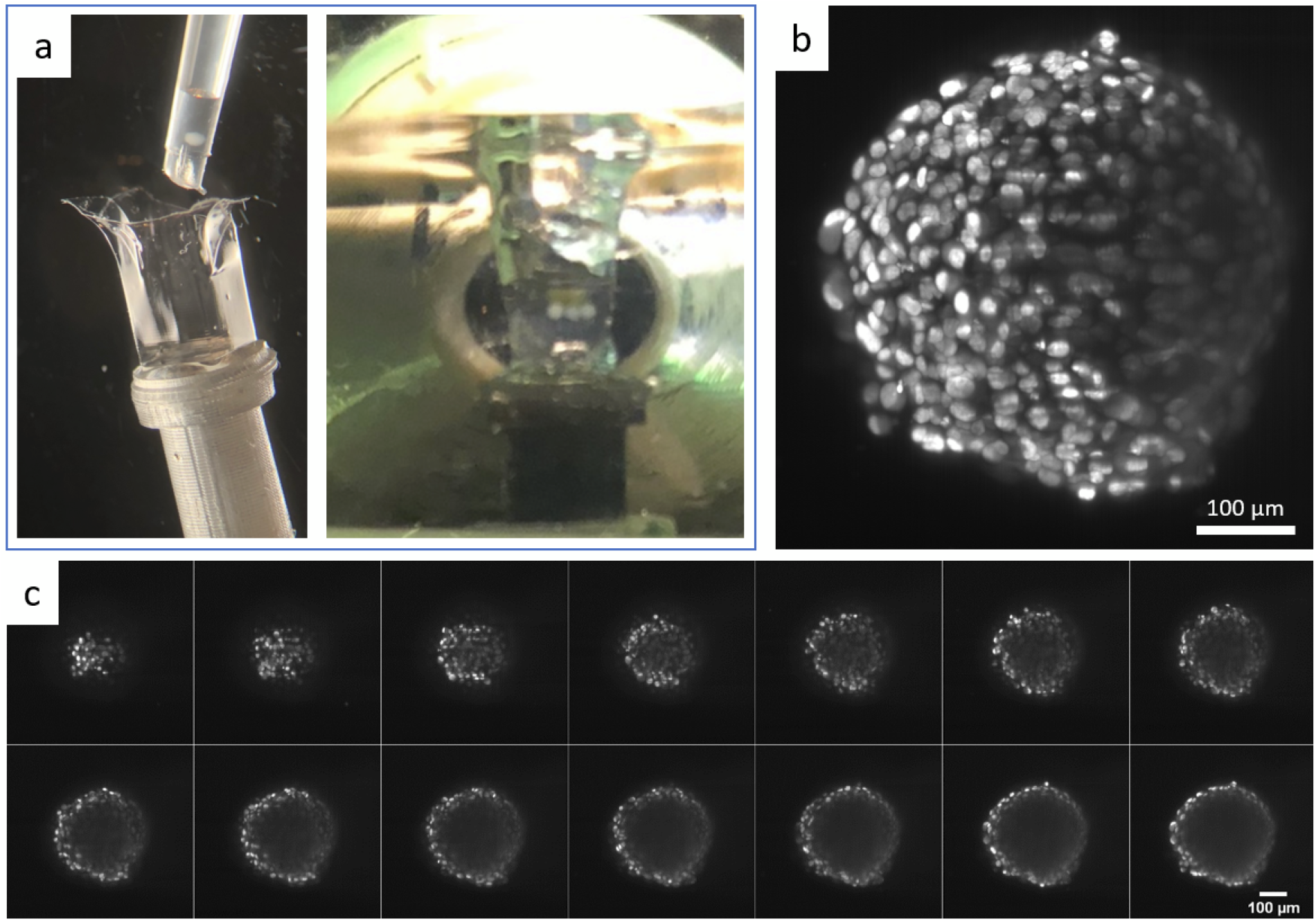
Imaging of cellular spheroids with the optical window. (a) Left: Human mesenchymal stem cell (hMSC) spheroids were pipetted in a FEP-foil cuvette for LSFM, as described in (14) and (15). Right: a FEP-foil cuvette containing three hMSC spheroids, visible as white adjacent dots in the image, was placed in the LSMF chamber in front of the objective lens covered with the optical window. (b) Maximum intensity projection of one hMSC spheroid from a stack comprising 200 slices. (c) Montage showing individual slices from the image stack of the same spheroid. Fluorescence staining: DAPI. Objective lens: Zeiss W N-Achroplan 10x/0.3 M27

In **Figure 9b**, a maximum projection of 200 individual slices is displayed. **Figure 9c** shows a montage of different planes of the same spheroid, with a z-distance of 10 µm between individual frames. As the spheroids were not optically cleared, the penetration depth is, as expected for compact spheroids, limited to the outermost cell layers.

### 6. Perfluorodecalin-compatible chambers for live honeybee imaging

A good example of the versatility and customizability of the proposed design methods for specimen chambers and sealing solutions is a chamber for imaging the development of live honeybee embryos over extended periods. Cultivating honeybee embryos for live imaging presents several unique challenges. The embryos must be maintained at a stable temperature of 34.5 °C. They cannot develop in water, requiring immersion in a highly oxygen-saturated liquid such as perfluorodecalin to ensure an adequate oxygenation. Perfluorodecalin is an expensive imaging medium that is prone to evaporation and can easily leak from even water-tight spaces due to its low surface tension and viscosity. Additionally, perfluorodecalin has a very low thermal mass, making it difficult to maintain a stable temperature through direct heating. A specimen chamber design that addresses all these challenges is shown in Figure 10. We reduced the volume of the specimen chamber to minimise the use and evaporation of perfluorodecalin; the heating element is installed in a separate water reservoir that provides thermal mass to stabilise the temperature; and all seals are lubricated with thick PTFE-based grease to prevent leakage. To the best of our knowledge, this design is the first to enable live honeybee embryo imaging in a light-sheet microscope.

**Figure 10.**
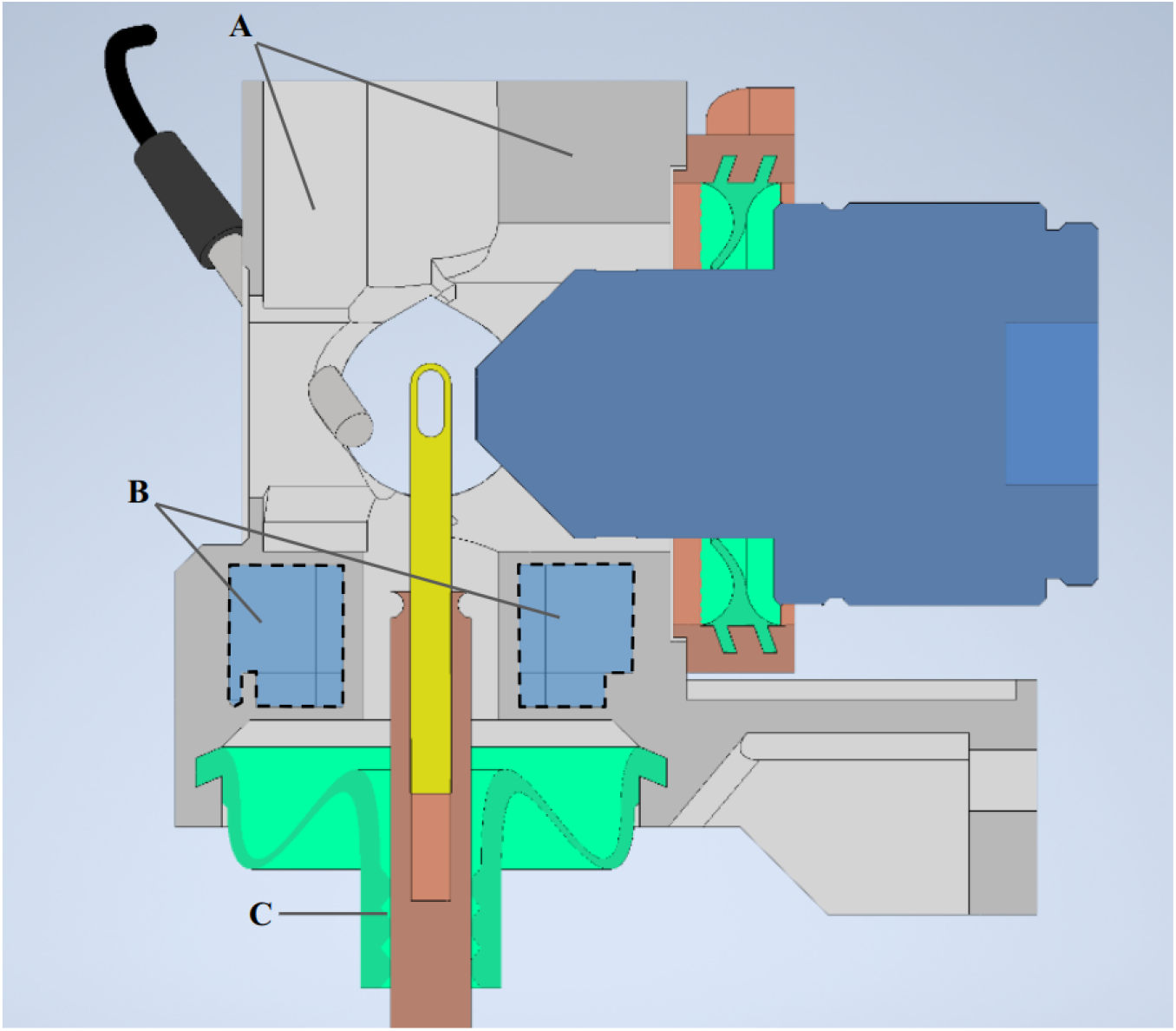
CAD model of a specimen chamber for live honeybee imaging. (A) volume reducing features, to minimize the amount of perfluorodecalin required for each imaging session. (B) separate compartment filled with water that contains a heating element installed directly in it. This setup provides substantial thermal mass, ensuring even heating throughout the chamber. (C) O-ring style features that are lubricated with PTFE-based grease, which is particularly effective at retaining perfluorodecalin within the chamber, despite the substance’s low viscosity and surface tension.

## Discussion

We presented open-hardware solutions for light sheet fluorescence microscopy (LSFM) chambers addressing longstanding and common challenges associated with the most widespread LSFM setups. By leveraging the versatility of 3D printing and innovative design approaches, we provide practical solutions that can be straightforwardly implemented at a laboratory scale with affordable equipment to enhance the functionality, user-friendliness, and adaptability of LSFM chambers.

One of the innovations presented in this work is the easy fabrication of highly flexible silicone connections and seals using a laboratory-scale injection moulding process. This approach allows the fabrication of thin, water-tight seals for objective lenses and bellows capable of accommodating large XYZ displacements and rotations of the sample holder. These components’ flexibility and low-force characteristics are critical for maintaining the precision of imaging objectives, particularly in setups requiring dynamic and fast focus adjustment. This innovation directly addresses the limitations of common and sub-optimal sealing solutions, which restrict the movement of the objective lenses.

Furthermore, the introduction of a thermoformed thin “optical window” between the imaging objective lens and the LSFM chamber interior offers a robust solution to the issues of contamination and chamber sterility. By isolating the objective lens from the immersion medium, this design not only prevents bacterial contamination during long-term imaging but also facilitates the rapid exchange of chambers and objectives without compromising the sterility of the experimental setup. Moreover, contamination or potential damage of the objective lens itself, especially when an organic clearing solution such as BABB is used, is also avoided. The use of fluorinated polymer (FEP) foil for the optical window, due to its favourable optical properties (high transparency, minimal thickness, refractive index close to the one of water) ensures minimal optical aberrations and good image quality.

The successful application of these chamber designs is showcased in a setup for the long-term live imaging of honeybee development and in the imaging of human mesenchymal stem cell spheroids. The demonstration of good-quality imaging using the optical window system, along with the possibility of maintaining chamber sterility during multiple imaging sessions, highlights the effectiveness of these technical solutions.

Overall, this study not only advances the technical capabilities of LSFM but also promotes accessibility and customization in microscopy by providing open-hardware systems that can be easily replicated and adapted to specific research needs. The open-source nature of the CAD and 3D printing files further democratises these advancements, enabling the broader scientific community to swiftly implement these solutions.

## Materials and Methods

### CAD modelling and design

CAD modelling of all parts except for optical window components was done in Autodesk Inventor. The optical window components were modelled in Autodesk Fusion 360. All 3D model files are available in the GitHub repository (16).

### Fabrication of specimen chambers and moulds for silicone injection moulding

All parts were 3D-printed using Bambu X1 Carbon FDM 3D-printer with 0.4 mm nozzle. STEP files after CAD modelling were sliced and prepared for printing using Bambu Studio slicer provided by our 3D-printer manufacturer. Precise settings used for slicing and printing can be found in Bambu studio project files in the GitHub repository (16).

Bambu Lab PLA Basic black filament was used for all the parts. Bambu Lab Support for PLA filament was used as an interface layer wherever parts required supports to be printed.

### Fabrication of seals via injection moulding of silicone

A two-component fast-curing dental silicone rubber eco-sil extrahart from picodent® was used to fabricate all silicone seals. The process for injection moulding was the following: clean and prepare the moulds; clamp them tightly together with nuts and bolts or built-in threaded joints; mix the two components silicone with about 10 ml extra over the expected injection volume; wait a bit and transfer to a separate container via a thin stream to minimise bubbles in the mixture; use a syringe without internal rubber seal to slowly suck up the silicone from the bottom of the container making sure to bleed the air bubble after the first several millilitres; hold the mould orienting it such that the air outlet is facing upward; slowly inject the silicone into the mould by pressing the syringe against the intake hole until you see silicone coming out from the air outlet; put the mould with the air outlet toward the bottom and plugged so that the silicone does not drip out; repeat with outer moulds. A video tutorial for injection moulding the silicone can be found at the GitHub repository (16).

All seals that require active movement of a part within the silicone seal, such as rotation of a shaft in a seal on Figure 6b,c, have been lubricated with Krytox GPL-205G0 PTFE grease to reduce friction.

### Fabrication of the optical window

The fabrication of the optical window using vacuum forming of FEP-foil (refer to (14) and (15)) was carried out by first cutting a 10 cm × 10 cm square patch of FEP-foil (50 μm thickness, Holscot Europe). The patch was then clamped into a small table-top vacuum-forming machine (JT-18, Jin Tai Machining Company), heated to 280 °C (surface temperature on the foil), and pressed onto the positive mould (**Figure 7**; the STL files are stored in the GitHub repository). Finally, the formed FEP optical window was cleaned with a detergent solution (1% Hellmanex-II in ultrapure water) and rinsed with abundant water.

Next, the optical window is cut to fit the open diaphragm for connection with the LSFM chamber. It is placed on the diaphragm and secured with a clamping ring (**Figure 8a**). The clamping ring is then glued to the adapter ring using either two-component epoxy glue or silicone rubber.

## Author contributions

A.G. and F.P. designed and manufactured the hardware. A.G. and F.P. wrote the manuscript. J.C. contributed to the design of the optical window, developed methods for thermoforming as well as prototyping with the two-component silicone rubber, and contributed to the writing of the manuscript. Support for research was provided by E.H.K.S. All authors discussed the results and edited the manuscript.

## Acknowledgements

We thank Sven Plath for technical assistance with the design of specimen chambers. A.G. F.P., and E.H.K.S. acknowledge funding by the Deutsche Forschungsgemeinschaft (DFG, German Research Foundation), grant GRK 2566 (project no. 414985841). F.P. acknowledges funding by the Wilhelm Sander-Stiftung (2020.008.1), the EU Horizon2020 project BRIGHTER (Grant #828931), the EU Horizon-EIC-2021 project B-BRIGHTER (Grant #101057894), and the German Space Agency at DLR (Grant #50W2019 and #50WB2316).

## Notes

### Competing Interest Statement

The authors have declared no competing interest.

## References

1. Huisken J, Swoger J, Del Bene F, Wittbrodt J, Stelzer EHK. Optical Sectioning Deep Inside Live Embryos by Selective Plane Illumination Microscopy. Science. 2004 Aug 13;305(5686):1007–9.

2. Keller PJ, Stelzer EH. Quantitative in vivo imaging of entire embryos with Digital Scanned Laser Light Sheet Fluorescence Microscopy. Curr Opin Neurobiol. 2008 Dec;18(6):624–32.

3. Stelzer EHK, Strobl F, Chang BJ, Preusser F, Preibisch S, McDole K, et al. Light sheet fluorescence microscopy. Nat Rev Methods Primer. 2021 Nov 3;1(1):73.

4. Krzic U, Gunther S, Saunders TE, Streichan SJ, Hufnagel L. Multiview light-sheet microscope for rapid in toto imaging. Nat Methods. 2012 Jul;9(7):730–3.

5. Schueth A, Hildebrand S, Samarska I, Sengupta S, Kiessling A, Herrler A, et al. Efficient 3D light-sheet imaging of very large-scale optically cleared human brain and prostate tissue samples. Commun Biol. 2023 Feb 13;6(1):1–15.

6. Diederich B, Lachmann R, Carlstedt S, Marsikova B, Wang H, Uwurukundo X, et al. A versatile and customizable low-cost 3D-printed open standard for microscopic imaging. Nat Commun. 2020 Nov 25;11(1):5979.

7. Lange M, Granados A, VijayKumar S, Bragantini J, Ancheta S, Santhosh S, et al. Zebrahub – Multimodal Zebrafish Developmental Atlas Reveals the State-Transition Dynamics of Late-Vertebrate Pluripotent Axial Progenitor [Internet]. 2023 [cited 2024 Aug 21]. Available from: 10.1101/2023.03.06.531398

8. Prem Ananth K, Jayram ND. A comprehensive review of 3D printing techniques for biomaterial-based scaffold fabrication in bone tissue engineering. Ann 3D Print Med. 2024 Feb 1;13:100141.

9. Shilov SY, Rozhkova YA, Markova LN, Tashkinov MA, Vindokurov IV, Silberschmidt VV. Biocompatibility of 3D-Printed PLA, PEEK and PETG: Adhesion of Bone Marrow and Peritoneal Lavage Cells. Polymers. 2022 Sep 22;14(19):3958.

10. Schmied C, Tomancak P. Sample Preparation and Mounting of Drosophila Embryos for Multiview Light Sheet Microscopy. In: Dahmann C, editor. Drosophila: Methods and Protocols [Internet]. New York, NY: Springer; 2016 [cited 2024 Aug 21]. p. 189–202. Available from: 10.1007/978-1-4939-6371-3_10

11. Pampaloni F, Berge U, Marmaras A, Horvath P, Kroschewski R, Stelzer EHK. Tissue-culture light sheet fluorescence microscopy (TC-LSFM) allows long-term imaging of three-dimensional cell cultures under controlled conditions. Integr Biol. 2014 Oct 22;6(10):988–98.

12. Keller PJ, Schmidt AD, Wittbrodt J, Stelzer EHK. Reconstruction of Zebrafish Early Embryonic Development by Scanned Light Sheet Microscopy. Science. 2008 Nov 14;322(5904):1065–9.

13. Kaufmann A, Mickoleit M, Weber M, Huisken J. Multilayer mounting enables long-term imaging of zebrafish development in a light sheet microscope. Development. 2012 Sep 1;139(17):3242–7.

14. Hötte K, Koch M, Hof L, Tuppi M, Moreth T, Verstegen MMA, et al. Ultra-thin fluorocarbon foils optimise multiscale imaging of three-dimensional native and optically cleared specimens. Sci Rep. 2019 Nov 21;9(1):17292.

15. Hof L, Moreth T, Koch M, Liebisch T, Kurtz M, Tarnick J, et al. Long-term live imaging and multiscale analysis identify heterogeneity and core principles of epithelial organoid morphogenesis. BMC Biol. 2021 Feb 24;19(1):37.

16. https://github.com/artgolden/light_sheet_specimen_chambers

